# Embodied schooling behavior via hydrodynamic interactions

**DOI:** 10.64898/2026.11.12.766162

**Authors:** Li-Ming Chao, Hun Jang, Máté Nagy, Laibing Jia, Amneet Pal Singh Bhalla, Boyce E. Griffith, Iain D. Couzin, Liang Li

**Affiliations:** Department of Collective Behaviour, Max Planck Institute of Animal Behavior, Konstanz 78464, Germany; Centre for the Advanced Study of Collective Behaviour, University of Konstanz, Konstanz 78464, Germany; Department of Biology, University of Konstanz, Konstanz 78464, Germany; Department of Engineering Mechanics, Inner Mongolia University of Technology, Hohhot 010051, China; Department of Biological Physics, ELTE Eötvös Loránd University, 1117 Budapest, Hungary; Institute of Evolution, HUN-REN Centre for Ecological Research, 1121 Budapest, Hungary; Department of Naval Architecture, Ocean and Marine Engineering, University of Strathclyde Glasgow G4 0LZ, UK; Department of Mechanical Engineering, San Diego State University, San Diego, CA 92182, USA; Departments of Mathematics, Applied Physical Sciences, and Biomedical Engineering, University of North Carolina, Chapel Hill, North Carolina 27599, USA; Carolina Center for Interdisciplinary Applied Mathematics, University of North Carolina, Chapel Hill, North Carolina 27599, USA; Computational Medicine Program, University of North Carolina School of Medicine, Chapel Hill, North Carolina 27599, USA; McAllister Heart Institute, University of North Carolina School of Medicine, Chapel Hill, North Carolina 27599, USA; Department of Computer and Information Science, University of Konstanz, 78464, Konstanz, Germany

## Abstract

Collective animal behavior is commonly explained by sensory-mediated interactions among individuals, yet physical interactions between moving bodies and their environment may also contribute to coordination. In fish schooling, passive hydrodynamic interactions have long been proposed to facilitate collective organization, but biological evidence remains limited because their effects are difficult to disentangle from active sensorimotor responses. Here, we combine computational simulations and physical experiments with self-propelled foils to identify balanced vortex interactions and develop a Vortex Balance Model (VBM) that predicts schooling configurations from swimming kinematics and relative tailbeat phase. We test these predictions using pairs of goldfish (*Carassius auratus*) swimming in a flow tank under three sensory conditions: intact vision and lateral-line sensing, visual deprivation with intact lateral-line sensing, and combined visual deprivation and reversible lateral-line impairment. Agreement between observed schooling configurations and VBM predictions increases as sensory input is reduced, reaching its highest level when both sensory modalities are impaired. These findings provide indirect biological evidence that passive hydrodynamic interactions contribute to fish schooling, while their effects can be modified or overridden by active sensorimotor responses.

## Introduction

Embodied intelligence emphasizes that intelligent behavior emerges from interactions between the body and its environment, offering a novel perspective on the integration of form, function, and control in biology, fluid mechanics, and robotics [1–4]. While traditionally applied to individual behavior, this concept also provides a framework for understanding group-level coordination [5–8].Collective movement is one of the most widespread forms of group coordination in nature, ranging from synchronized fish schools [9–14] and coordinated bird flocks [15–18] to sheep herds [19] and dynamic human crowds [20–22]. These widespread patterns have attracted growing interest across biology, physics, and engineering [23] and are commonly attributed to active interactions, in which individuals use sensory information to adjust their behavior in response to neighbors or environmental cues [24–28].

Although fish schooling is commonly understood as a process driven by sensory perception and behavioral responses to neighbors, it has long been proposed that interactions mediated by the surrounding flow may also contribute to collective coordination [29, 30]. This possibility has attracted considerable interest in physics and engineering, where computational and robotic studies have demonstrated that coordinated swimming can emerge through hydrodynamic interactions alone [31–35]. However, whether such interactions contribute to the schooling behavior of living fish remains unclear. In natural fish schools, hydrodynamic interactions occur alongside continuous sensory feedback and behavioral adjustments, making it difficult to distinguish the contribution of the surrounding flow from that of active control. Consequently, direct biological evidence for hydrodynamically mediated schooling remains limited.

Here, we investigate whether passive hydrodynamic interactions contribute to schooling behavior in living fish. We combined computational fluid dynamics (CFD) simulations and experiments with bio-inspired foils to examine how interactions through the surrounding flow can give rise to coordinated swimming without sensory feedback or active behavioral control. Based on the resulting flow patterns, we developed a simplified Vortex Balance Model (VBM) that predicts the relative positions of swimmers from their kinematics alone, without requiring direct measurements of the surrounding flow. We then tested these predictions in freely swimming goldfish (*Carassius auratus*) under different sensory conditions, selectively reducing visual and lateral-line input to examine whether the contribution of passive hydrodynamic interactions becomes more apparent when sensory information is limited. Model predictions agreed more closely with observed fish behavior as sensory input was reduced. These findings suggest that passive hydrodynamic interactions can contribute to fish schooling, although their influence may be modified or overridden by active sensorimotor control.

## Results

### Computational fluid dynamics simulations

We investigated passive hydrodynamic interactions by simulating two identical, self-propelled pitching foils (Figure 1A), each free to translate with two-degrees-of-freedom (2-DOF) in the streamwise (*u*) and lateral (*v*) directions. Each foil had a semicircular leading edge, a body length *BL*, and a maximum thickness of 1*/*6 *BL* [36]. We resolved the surrounding flow and fluid–structure interactions using an adaptive-mesh immersed boundary method [37] (Figures 1B, S1 and S2; Text S1).

**Fig 1.**
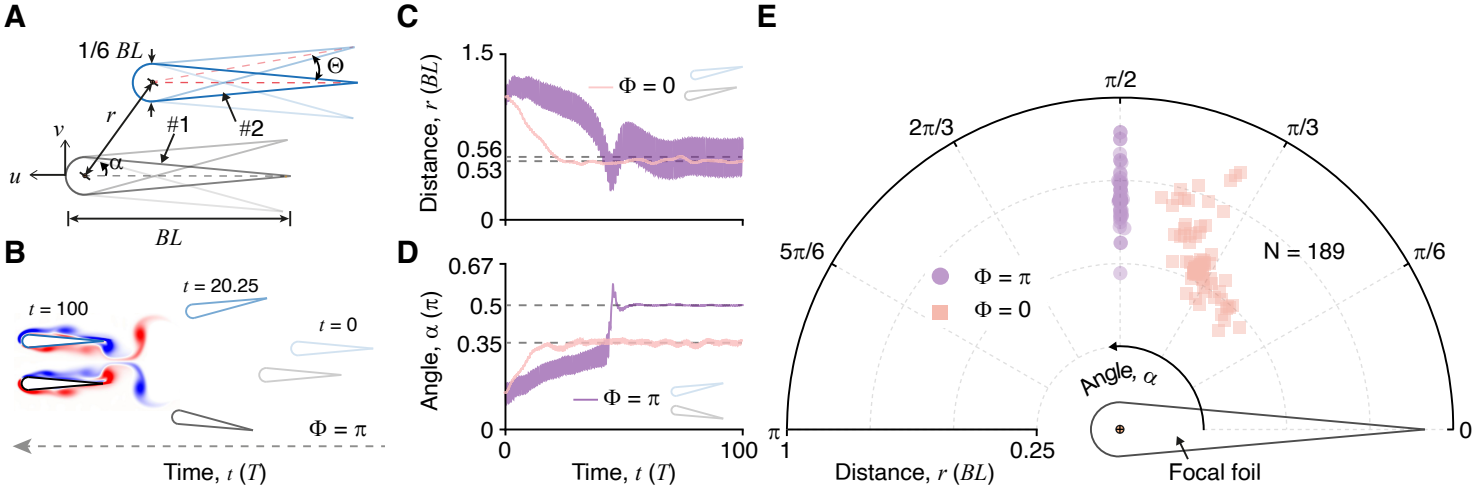
Embodied schooling observed in computational fluid dynamics simulations. (A) Geometric configuration of two identical foils (#1 and #2). Each foil had a semicircular leading edge, a body length *BL*, and a maximum thickness of 1*/*6 *BL*. Both foils are free to move in the streamwise (*u*) and lateral (*v*) directions. The pitching amplitude is denoted as Θ. The relative distance *r* and stagger angle *α* between the two foils are defined in a polar coordinate system based on the pitching centers. (B) Representative flow patterns observed in simulations at selected nondimensionalized times *t*, where *t* is normalized by the pitching period *T*. (C, D) Examples of time evolution of *r* and *α* at Φ = *π* (out-of-phase condition) and Φ = 0 (in-phase condition), respectively, with dynamic schooling patterns indicated. (E) Time-averaged relative positions of foil #2 on the (*r, α*) map relative to focal foil #1. Two types of time-averaged relative configurations emerged, determined by the initial phase difference Φ: Φ = *π* results in a side-by-side arrangement and Φ = 0 causes a staggered formation, regardless of the initial spatiotemporal conditions and flow parameters. Each marker represents the final time-averaged relative position of foil #2 obtained from an independent trial with slightly different initial conditions. The algorithm used to identify the equilibrium positions from the CFD data is presented in Algorithm S1.

To systematically investigate embodied hydrodynamic interactions in 2-DOF, we performed CFD simulations spanning pitching period (*T* ∈ [0.4, 1.0] s), pitching amplitude (Θ ∈ [0.056, 0.139] *π*), phase difference (Φ, with Φ = 0 for in-phase and Φ = *π* for out-of-phase), center-to-center distance (*r*, with 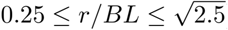), stagger angle (*α*, with 0 *≤ α/π ≤* 0.5) (Figures 1A and S3), and fluid viscosity (*µ* ∈ [0.0005, 0.005] poise). Among these, 189 cases developed embodied schooling: 95 exhibited an oscillatory side-by-side pattern with Φ = *π*, and 94 an oscillatory staggered pattern with Φ = 0 (Figures 1C-1E; Video S1). The remaining cases resulted in collision or separation (Figure S4). By randomly pairing trajectories across all 600 tested cases, we found that only 0.7% (Figures S5-S7; Video S2) exhibited schooling behavior, significantly lower than the 31.5% observed in the CFD ensemble (*P* = 0.001, Figure S5). Further analyses showed that greater initial distances led to longer convergence times, but yielded similar schooling patterns across the tested pitching frequencies and amplitudes (Figures S8 and S9). Moreover, the side-by-side schooling pattern exhibited smaller fluctuations in stagger angle *α* but larger variations in distance *r*; whereas the staggered pattern showed comparable fluctuations in both *r* and *α*, with slightly greater overall variability (Figures 1C, 1D and S10-S13). We also observed that when the two foils operate out of phase, they predominantly maintain forward motion along the streamwise direction. In contrast, in-phase motion resulted in pronounced lateral displacement of the pair (Figure S14).

### Experiments with pairwise flapping foils

We further experimentally investigated the embodied hydrodynamic interactions of pairwise flapping foils in a flow tank (Loligo system, Tjele, Denmark). Guided by the CFD results (Figure S15), we conducted representative experiments at two Reynolds numbers (*Re*) by varying the flow speed (Figure 2A). Instead of allowing the foils to self-propel, we measured the resulting forces in both the streamwise and lateral directions using high-resolution six-axis force sensors (K6D40 50N/5Nm/MP11, ME-Meßsysteme GmbH) mounted on each flapping foil (Figures 2B and S16; Video S3; Text S2). The prescribed conditions were selected based on isolated-foil measurements showing approximately zero time-averaged net streamwise force, thereby approximating self-propelled swimming [38]. The experiments tested the initial relative position between the two foils in Cartesian coordinates, defined as (Δ_FB_, Δ_LR_), with body length *BL* = 100 mm. Here, Δ_FB_ denotes the front-back offset between the two foils and Δ_LR_ denotes the left-right spacing. For in-phase motion (Φ = 0), the tested positions covered Δ_FB_ ∈ [*−*10, 100] mm with Δ_FB,step_ = 10 mm, and Δ_LR_ ∈ [80, 110] mm with Δ_LR,step_ = 10 mm, corresponding to initial distances of 0.8 to 1.49 *BL*. For out-of-phase motion (Φ = *π*), the tests focused on side-by-side configurations with Δ_FB_ ∈ [*−*10, 10] mm and Δ_FB,step_ = 5 mm, and Δ_LR_ ∈ [80, 110] mm, where the tested Δ_LR_ values were spaced by 5-10 mm, corresponding to initial distances of 0.8 to 1.1 *BL*. The flow speeds were 0.108 and 0.15 m*/*s, equivalent to 1.08 and 1.5 *BL/s*. The pitching periods ranged from 0.5 to 1.0 s, the pitching amplitudes ranged from 0.069*π* to 0.125*π*, and the phase differences were Φ = 0 and Φ = *π*.

**Fig 2.**
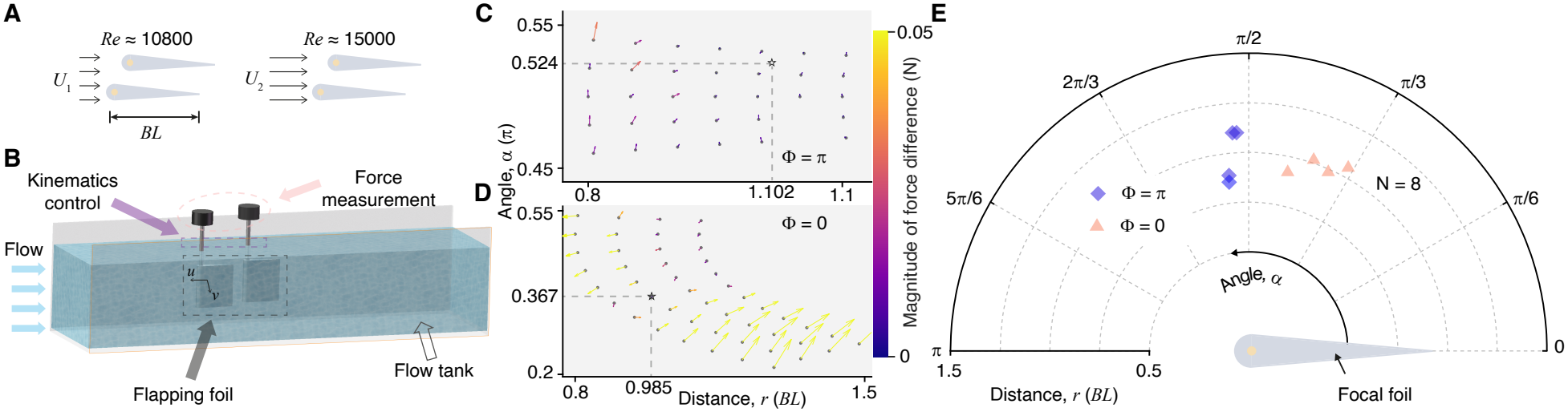
Hydrodynamic interactions between two flapping foils in a flow tank. (A) Representative experiments conducted at two Reynolds numbers (*Re*). (B) Experimental platform featuring high-resolution six-axis force sensors mounted on each flapping foil to measure forces in both the streamwise (*u*) and lateral (*v*) directions. (C, D) Force-imbalance vectors (arrows) between the two foils plotted on the (*r, α*) map. Zero-magnitude positions were identified based on the directions and magnitudes of the force vectors, in which the force differences between the two foils vanish in both the streamwise and lateral directions; these positions are interpreted as candidate force-balanced schooling configurations. The algorithm used to identify the zero-magnitude positions from the experimental data is presented in Algorithm S2. (E) Observed side-by-side and staggered swimming configurations across experiments with varying relative angles and phase differences.

Because the experimental foils were held fixed during force measurements, we used the cycle-averaged force differences between the two foils to identify candidate schooling configurations (Figures 2C and 2D; Text S2). Across 32 tests, we identified eight candidate schooling configurations in which the measured mean force differences vanished in both the streamwise and lateral directions. Quasi-side-by-side and staggered configurations were observed for Φ = *π* and Φ = 0, respectively (Figures 2E). These results suggest that passive hydrodynamic coupling can produce similar schooling patterns across a broad range of Reynolds numbers and flapping kinematics [39] (Figures S17 and S18).

### Mechanism of embodied hydrodynamic interactions in schooling behavior

To examine the hydrodynamic mechanism underlying embodied schooling, we first conducted flow visualizations in both simulations and experiments with two foils (Figure S19). In both settings, we observed coherent vortex interactions associated with out-of-phase side-by-side schooling (Figure 3A) and in-phase staggered schooling (Figure 3B). These observations motivated a simplified Vortex Balance Model (VBM), in which the geometry of the shed reverse Bénard–von Kármán vortices determines the balance of induced velocities. The resulting predictions for schooling distance and stagger angle are given by Equation 1, with the derivation provided in Text S3.

**Fig 3.**
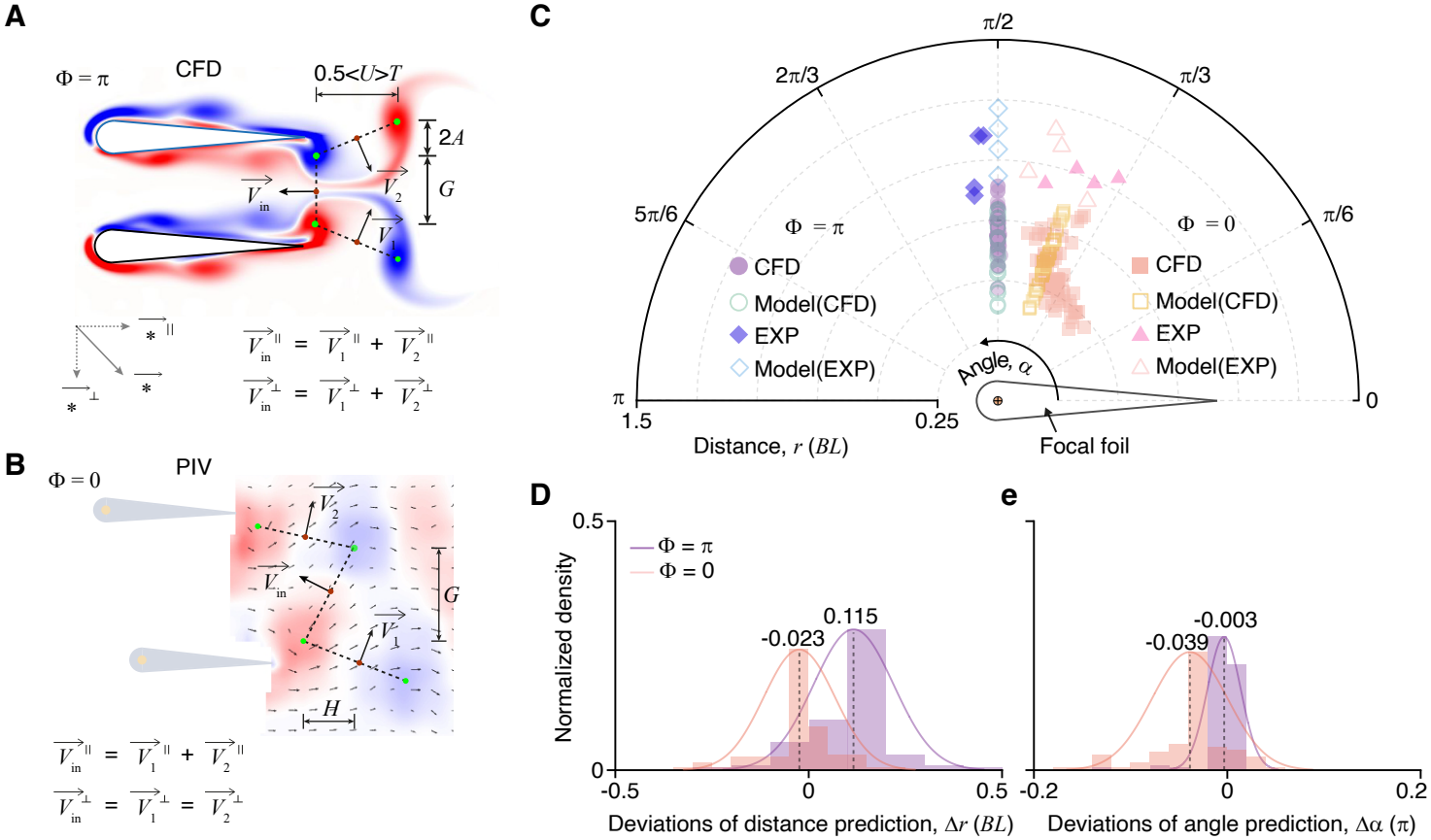
Vortex Balance Model (VBM). (A, B) Schematics of two swimmers under out-of-phase (Φ = *π*) and in-phase (Φ = 0) conditions. The flow structures generated by free-swimming bodies are idealized as reverse Bénard–von Kármán vortex streets. The streamwise, *H* and lateral, *G* spacings between adjacent vortices are estimated as 0.5 ⟨*U T*⟩ and 2*A*, where ⟨*U*⟩ is the time-averaged swimming speed and *A* is the tailbeat amplitude (Figures S19 and S20). When embodied schooling is achieved, the induced velocity components satisfy: 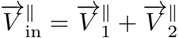 and 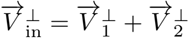 for Φ = *π*, and 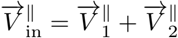 and 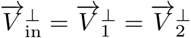 for Φ = 0. Here, 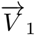 and 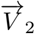 denote the velocities induced by the dipoles shed by swimmers #1 and #2, and 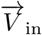 is the velocity induced by the inner-side vortex pair. Superscripts ^∥^ and ^⊥^ refer to the longitudinal and lateral components of the velocity vectors. (C) Comparison of model predictions with extended CFD simulations and experiments with foils (EXP), using kinematic parameters measured from the CFD/EXP studies. (D, E) Deviations of distance and angle between CFD/EXP data and model predictions.

In Equation 1, ⟨*U*⟩ denotes the average swimming speed, *A* refers to tailbeat amplitude (Figure S20), and 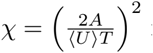 is the square of the Strouhal number. Using VBM, we evaluated all cases in which embodied schooling (*r, α*) emerged purely through hydrodynamic interactions, based on kinematic parameters (*T, A*, ⟨*U*⟩) and phase difference (Φ). The results show that the model performs well for both simulations and experiments, with deviations of approximately 0.039 *π* in *α* and 0.023 *BL* in *r* for Φ = 0, and 0.003 *π* in *α* and 0.115 *BL* in *r* for Φ = *π* (Figure 3D and 3E). These deviations may partly reflect the idealized, inviscid representation of the vortex structures [40], comparison between the present VBM predictions and previously reported schooling configurations has been shown in Figure S21.

### VBM verification with real fish

To assess whether passive hydrodynamic interactions contribute to fish schooling behavior, we conducted experiments with freely swimming pairs of goldfish in a flow tank under conditions of intact and reduced sensory input. Specifically, we manipulated the availability of visual and lateral-line information by testing fish either under visible illumination or under infrared illumination outside their visible spectrum, with lateral-line function either intact or reversibly impaired, across a range of flow speeds (see Methods for details). This resulted in four treatment groups: fully functional (V+LL+), vision only (V+LL-), lateral line

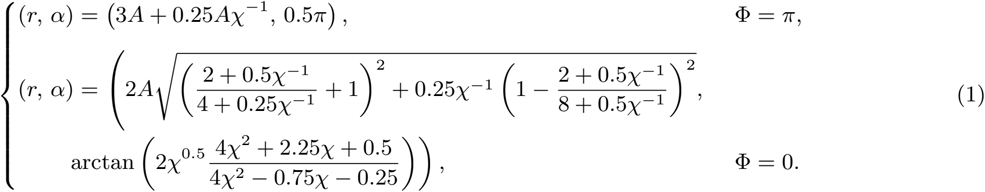

only (V-LL+), and both systems impaired (V-LL-). Using deep learning-based tracking to extract kinematic data, we estimated the potential swimming configurations for both in-phase and out-of-phase swimming. A comparison between the real fish relative positions and the schooling patterns predicted by the VBM is shown in Figure 4. As sensory input was reduced, the observed schooling distributions became more concentrated (Figures 4A, 4D, 4G and 4J). When kinematic parameters were used as input to the VBM (Figures 4B, 4E, 4H and 4K), the deviations in predicted distances and angles were similarly uniformly distributed, with small and comparable energy distances (a statistical measure of dissimilarity between two probability distributions based on pairwise distances [41]) across conditions (V+LL+ vs V-LL+: 0.17; V+LL+ vs V+LL-: 0.19; V-LL+ vs V+LL-: 0.06). In contrast, the joint prediction-error distribution for V-LL-differed more strongly from those of the other sensory conditions and from all randomized controls, with energy distances of 0.39 (vs V+LL-, *P* = 0.001), 0.41 (vs V-LL+, *P* = 0.001), and 0.53 (vs V+LL+, *P* = 0.001) (Figure 4M).

**Fig 4.**
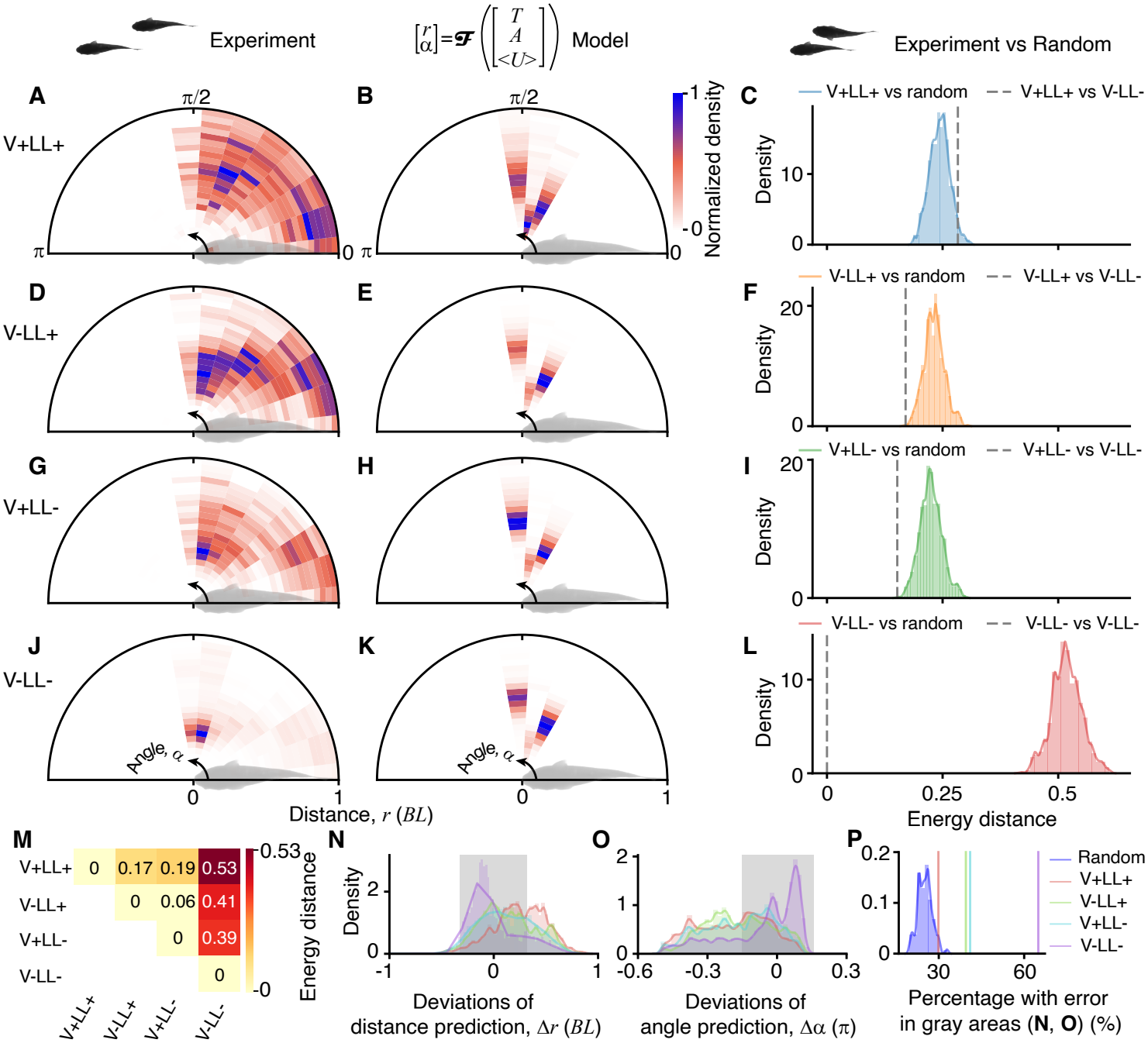
Vortex Balance Model (VBM) verification with real fish. (A-L) Spatial distributions of neighboring fish (A, D, G, J) and theoretical predictions of schooling patterns from the Vortex Balance Model (VBM) (B, E, H, K), shown relative to the focal fish during retained interaction episodes classified as either in-phase or out-of-phase swimming across all tested conditions: (A, B) both visual and lateral line systems enabled (V+LL+); (D, E) lateral line only (V-LL+); (G, H) vision only (V+LL-); and (J, K) both systems impaired (V-LL-). (C, F, I, L), Energy distances quantifying prediction errors between random pairs and V+LL+ (C), V-LL+ (F), V+LL-(I), and V-LL-(L), where the distributions represent energy distances from the corresponding conditions to random pairs and dashed lines indicate energy distances from V-LL- to all other conditions. (M-P) Energy distances between each pair of conditions (M); deviations between prediction and data in distance (N, gray area) within 0.4 *BL* and angle (O, gray area) within 0.3 rad for all conditions; and percentage of prediction errors within the gray area in (N) and (O) for all conditions and random pairs (P).

We further examined randomized pairings of fish trajectories under the V-LL-condition (Figure S22; Video S4) to examine whether the differences could be attributed to random interactions. The results showed that random pairs more closely resembled V+LL+, V-LL+, and V+LL-conditions (the distributions in Figures 4C, 4F, 4I and 4L), with energy distances smaller than 0.1, while all random cases exhibited larger energy distances compared to V-LL-(the dashed line in Figures 4C, 4F, 4I and 4L). These results indicate that V-LL-is significantly different from all other conditions, and that the other three conditions are not significantly different from random pairs. Moreover, when considering the percentage of data points within a 0.4 *BL* deviation and 0.3 rad angular deviation between the data and prediction (gray areas in Figures 4N and 4O), we observe that 64.97% of cases in the V-LL-condition met this criterion. This percentage was significantly higher (*P* = 0.001) than those in other conditions: 29.86% for V+LL+, 39.52% for V-LL+, 41.01% for V+LL-, and an average of 25.11% for randomly paired controls (standard error: 2.46%, Figure 4P). Together, these results suggest that passive hydrodynamic interactions contribute to fish schooling and become more apparent when sensory input is reduced. When sensory information is available, active sensorimotor control can readily override these lower-level embodied hydrodynamic interactions.

## Materials and methods

### Numerical methodology

Flapping motions used in numerical simulations are defined as:

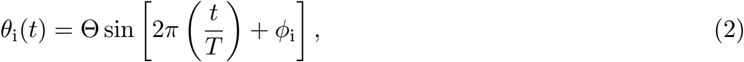

where *i* denotes the foil index, Θ is the maximum pitching amplitude, *t* is time, *T* is the pitching period, and *ϕ*_*i*_ is the initial phase of foil *i*. To account for the two possible initial directions of foil motion, the initial phase of foil #1 was set to either *ϕ*_1_ = 0 or *ϕ*_1_ = *π*. For each value of *ϕ*_1_, the phase of foil #2 was prescribed such that the relative phase difference Φ = *ϕ*_2_ *− ϕ*_1_ was either 0 or *π*. Specifically, the in-phase condition (Φ = 0) comprised (*ϕ*_1_, *ϕ*_2_) = (0, 0) and (*π, π*), whereas the out-of-phase condition (Φ = *π*) comprised (*ϕ*_1_, *ϕ*_2_) = (0, *π*) and (*π*, 0). These two relative phase relationships were considered because they are commonly observed during collective swimming [9].

The numerical simulations employed the IBLevelSet method implemented in the open-source IBAMR software [37]. IBAMR is a distributed-memory parallel implementation of the immersed boundary (IB) method [42, 43] that integrates the Cartesian grid adaptive mesh refinement (AMR) technique. IBAMR has been widely employed in studies of fish-like propulsion [44–46]. More details about numerical methodology can be found in Text S1.

In simulations, we first considered a specific combination of parameters, e.g., *T* = 1 s, Θ = 0.111 *π* and *µ* = 0.001 poise. The initial spatial separation between the two foils was configured using a hybrid grid sampling strategy, as illustrated in Figure S3. A refined rectangular grid with a resolution of 0.25 *×* 0.1 *BL* was employed for the domain 0 *≤ FB*_0_ *≤* 1 *BL* and 0.5 *≤ LR*_0_ *≤* 1 *BL*. In addition, a coarser grid with 0.25 *×* 0.25 *BL* resolution was applied to two regions: (i) 1.25 *≤ FB*_0_ *≤* 1.5 *BL* and 0 *≤ LR*_0_ *≤* 1 *BL*, and (ii) 0 *≤ FB*_0_ 1.5 *BL* and 1.25 *≤ LR*_0_ *≤* 1.5 *≤ BL*. Here, *FB*_0_ and *LR*_0_ denote the initial front–back and left–right separations of foil #2 relative to foil #1, respectively. Numerical simulations under this parameter combination indicated that the resulting schooling patterns were insensitive to the initial spatial separation when the pitching frequency, amplitude, and fluid viscosity are fixed (Figures S8 and S9). Therefore, for simulations involving other combinations of flapping kinematics and fluid properties, we considered only four different initial positions, selected using Latin hypercube sampling.

The simulations spanned a range of flapping frequencies (1.0, 1.5, 2.0, and 2.5 Hz), prescribed input amplitudes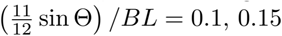, 0.2, and 0.25, phase differences Φ = 0 and *π*, and fluid viscosities *µ* = 0.0005, 0.001, and 0.005 poise. The results of grid independence tests are shown in Figure S2. All simulations consumed at least 2.5 million core hours in the present work. Data analyses of numerical results are detailed in Text S4.

### Flapping foil experimental setup

Flow tank experiments were conducted using two flapping foils mounted on the RoboTwin platform [47] at the Max Planck Institute of Animal Behavior, Konstanz, Germany. The RoboTwin platform consisted of a flow tank (Loligo Systems, Tjele, Denmark), a high-resolution recording and tracking system, a six-axis motion platform, and an integrated system for measuring power consumption and thrust generation. Each foil was designed with a two-dimensional teardrop-shaped profile, closely resembling the geometry used in simulations. The chord length (*BL*) was 100 mm (Figure 2A), and the span length was 120 mm. Pitching motion was actuated by DYNAMIXEL XL430 servo motors.

For force measurements, each foil was mounted on an ME K6D40 six-axis force sensor, which recorded both streamwise (thrust) and lateral (lift) forces. Over the course of the experiments, more than 2,500 individual cases were systematically measured and analyzed. All two-foil experiments were initialized from self-propelled conditions identified in isolated-foil experiments, corresponding to approximately zero net mean force. The force measurements were validated using three repeated trials (Figure S23). A force-balanced configuration was defined as one in which the two foils experienced equal time-averaged thrust and lift forces. The experiments were performed across a range of flapping frequencies (1.0, 1.5, and 2.0 Hz), normalized prescribed input amplitudes (0.069, 0.097, and 0.125 *π*), and phase differences (Φ = 0 and *π*). Furthermore, under the present experimental conditions, the measured hydrodynamic forces of two flapping foils were dominated by the inter-foil interactions, while the available boundary-effect tests indicated limited sensitivity to lateral confinement (Figures 3B, S24 and S25; Text S2). A visual summary of the flow tank experiments is provided in Video S3. Two-dimensional particle image velocimetry (PIV) measurements were also performed using this platform (Text S5).

### Vortex Balance Model (VBM)

Motivated by the vortex interactions observed during embodied schooling in simulations and experiments (Figure 3A, 3B and S19; Video S5), we modeled the balance of velocities induced by reverse Bénard–von Kármán vortex streets. For the out-of-phase case (Figure 3A), the induced velocity components satisfy 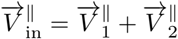 and 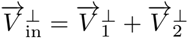. For the in-phase case (Figure 3B), the two sets of vortex streets are spatially shifted, and the balance conditions are 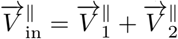 and 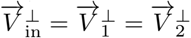. Here, 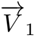 and 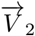 denote the velocities induced by the reverse Bénard–von Kármán dipoles shed by swimmers #1 and #2, respectively, and 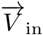 represents the velocity induced by the two inner-side vortices within the wake. Superscripts ^∥^ and ^⊥^ indicate the streamwise and lateral components of the induced velocity vectors. These assumptions yield analytical expressions for the streamwise component *H* and lateral component *G* associated with the assumed vortex balance (Figure 3A and 3B):

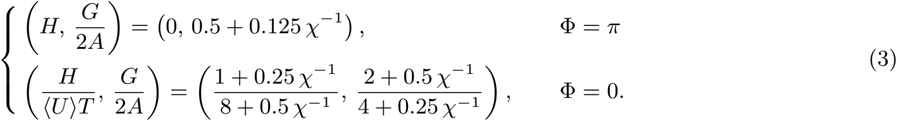

Thus, the corresponding schooling patterns are given by (*r, α*) = (*G* + 2*A*, 0.5*π*) when Φ = *π* and 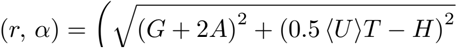 , arctan 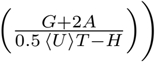 when Φ = 0, respectively. Further details on the derivation are provided in the Text S3.

### Fish experiments

Fish experiments were conducted with freely swimming pairs of goldfish using the RoboTwin platform in a recirculating flow tank. The effective test area of the tank was 250 mm wide, 875 mm long, and 250 mm deep. A mirror was positioned at the bottom of the tank at 45° to the horizontal plane, allowing a camera mounted beside the tank to record the reflected bottom view, while a second camera recorded the direct side view (Figure S26). The two views were synchronized and recorded using BASLER acA2000-165umNIR cameras at 100 frames per second with a resolution of 2048 *×* 1088 pixels. Videos were acquired using commercial recording software provided by Loopbio (Austria).

For each experimental session, two fish were selected and transferred to an acclimation container that was gradually filled with water from the flow tank. After 30 min, and once the temperature difference between the acclimation container and the flow tank was below 2 ^°^C, the fish were transferred to the flow tank and allowed to acclimate for a further 30 min without flow. During this period, calibrated images of each fish were acquired to determine body length. The sensory manipulations followed the between-pair design described under Experimental Model and Study Participant Details; fish were not reassigned between treatment groups.

Before each experiment, the flow speed was calibrated using a vane-wheel flow probe (Höntzsch, Germany) and adjusted according to the mean body length of the pair. Each pair was tested at five flow speeds ranging from 1.2 to 1.6 *BL* s^−1^ in increments of 0.1 *BL* s^−1^. The flow speeds were presented in randomized order.

Each speed treatment consisted of a 5-min recording period followed by a 5-min rest period without flow. White visible-spectrum illumination was used in the V+LL+ and V+LL-conditions. Infrared illumination at 850 nm was used in the V-LL+ and V-LL-conditions to substantially reduce visual input while allowing video recording with near-infrared-sensitive cameras. Lateral-line function was reversibly impaired following the previously established protocol [6]. Fish were immersed for 3 h in 6 L of water containing 2.7 g of neomycin, corresponding to a concentration of 0.45 g L^−1^. Following exposure, fish were tested using the same acclimation and recording procedures as fish with intact lateral-line function. Treatment efficacy was evaluated after the experiments using DASPEI (2-[4-(dimethylamino)styryl]-1-ethylpyridinium) staining. A randomly selected treated fish was immersed in a 0.006% DASPEI solution for 10 min and was subsequently anesthetized by immersion in 0.15 g L^−1^ MS-222. The lateral-line neuromasts were then examined using an epi-fluorescence stereomicroscope (ApoTome.2). Lateral-line impairment was considered successful when the neuromasts exhibited no visible DASPEI fluorescence, indicating a lack of dye uptake by the lateral-line hair cells. By contrast, the neuromasts of an untreated control fish exhibited clear fluorescence (see our previous work [6] for details). Fish positions, body postures, and swimming kinematics were extracted from the synchronized recordings as detailed in Text S6 and Figures S27-S29.

All experiments were conducted in accordance with the animal ethics permit approved by Regierungspräsidium Freiburg, 35-9185.81/G-17/90.

## Discussion

Our results provide biological evidence consistent with a contribution of passive hydrodynamic interactions to fish schooling, particularly when sensory input is reduced. By combining CFD simulations, physical experiments, and a simplified Vortex Balance Model (VBM), we identified dynamic schooling patterns could arise from passive hydrodynamic interactions. We further tested this mechanism in living fish and found that VBM predictions became more accurate as sensory input was reduced. These findings suggest that sensory-mediated responses can dominate schooling behavior, potentially masking the underlying contributions of passive hydrodynamic interactions. This may explain why direct biological evidence for this long-standing hypothesis has remained difficult to obtain, despite extensive theoretical investigations and the identification of plausible physical mechanisms.

We also noticed that discrepancies between observed fish behavior and VBM predictions persisted even when both the lateral line and vision were impaired (Figure 4J). These differences may reflect additional mechanisms involved in swimming coordination, including proprioceptive feedback [48], local reflexive responses [49], or changes in the surrounding flow induced by flexible caudal-fin movements [50]. These observations suggest that embodied hydrodynamic interactions may contribute to fish schooling alongside sensory and motor processes, rather than operating independently of them [28].

Fish schools are inherently dynamic and three-dimensional, with individuals continuously adjusting their relative positions rather than maintaining fixed arrangements over extended periods [51]. Similar dynamical behavior has also been discussed in recent experiments with flapping foils [52]. The schooling patterns identified here should therefore be understood as dynamic configurations that may emerge transiently within an evolving group, rather than fixed spatial arrangements. In the CFD simulations and foil experiments, these patterns arose through hydrodynamic interactions without explicit communication or sensory control. The live-fish experiments further suggest that similar interactions may contribute to schooling during natural body undulations and caudal-fin propulsion, although fish swimming involves more complex three-dimensional flows and remains coupled to sensorimotor control [53]. These results suggest that embodied hydrodynamic interactions may contribute a low-level physical mechanism for collective organization, upon which additional sensory and behavioral processes act in natural fish schools.

Taken together, our findings reveal a previously underappreciated contribution of passive hydrodynamic interactions to fish schooling and establish a tractable framework for investigating embodied intelligence in collective animal behavior. Beyond advancing our understanding of how fish coordinate their movements, the VBM also offers potential design principles for underwater swarm robotics, where fluid-mediated interactions could complement active sensing and control.

## Supporting information

S1 Fig. Adaptive mesh refinement for the CFD simulations. The four Cartesian levels resolve the moving foils and their wakes. Vorticity-magnitude thresholds of 0.25, 0.5, 1, and 2 s^−1^ were used on levels 0-3, respectively.

S2 Fig. Grid convergence for isolated and interacting foils. Three grid systems were considered: Grid system 1 (GS1), Δ*x* = Δ*y* = 6 *×* 10^−3^*BL*; GS2, 1.25 *×* 10^−3^*BL*; and GS3, 0.8 *×* 10^−3^*BL*. (A) Swimming speed of an isolated foil at (*f*, Θ, *µ*) = (1 Hz, 0.111*π*, 0.001 poise). (B,C) Relative distance *r* and stagger angle *α* for in-phase and out-of-phase foil pairs, respectively, initialized at (*r, α*) = (1.097*BL*, 0.260*π*). GS2 and GS3 yield closely matching trajectories, supporting the use of GS2.

S3 Fig. Initial-position sampling in the two-foil CFD simulations. Positions of foil #2 are shown relative to focal foil #1. The front-back and lateral offsets determine the initial distance and stagger angle; the sampling is denser at smaller separations.

S4 Fig. Representative outcomes from different initial positions. (A,B) Out-of-phase (Φ = *π*) and in-phase (Φ = 0) simulations, respectively, at (*f*, Θ, *µ*) = (1 Hz, 0.111*π*, 0.001 poise). Initial positions can lead to embodied schooling or collision, illustrating the dependence of the outcome on starting geometry.

S5 Fig. Randomized-pairing control for the CFD simulations. The distribution of the fraction of randomized trajectory pairs classified as schooling is compared with the fraction in the original interacting simulations. Schooling occurs more frequently in the interacting ensemble (*P* = 0.001).

S6 Fig. Randomized trajectory pairs classified as schooling. (A) Relative distance and (B) stagger angle for five pairs of trajectories drawn from separate simulations. These examples demonstrate that occasional schooling-like relative trajectories can arise without direct simulated interaction between the paired foils.

S7 Fig. Randomized trajectory pairs not classified as schooling. (A) Relative distance and (B) stagger angle for 251 pairings exhibiting separation or collision. Colors distinguish individual pairings and do not represent a physical variable.

S8 Fig. Influence of initial positions on the final mean schooling geometry. (A,B) Deviations of distance *r* from its mean across initial positions for out-of-phase and in-phase cases, respectively. (C,D) Corresponding deviations of stagger angle *α*. The narrow distributions indicate similar mean configurations among the cases that develop schooling.

S9 Fig. Evolution of relative distance and stagger angle from different initial positions. (A,B) Out-of-phase (Φ = *π*) cases; (C,D) in-phase (Φ = 0) cases. Each curve represents a separate CFD simulation at (*f*, Θ, *µ*) = (1 Hz, 0.111*π*, 0.001 poise). The trajectories illustrate different transients approaching similar schooling geometry.

S10 Fig. Out-of-phase schooling distance across kinematic and viscosity conditions. Time histories of relative distance *r* for Φ = *π* under the pitching frequencies *f* , amplitudes Θ, and viscosities *µ* indicated in each panel. Oscillation amplitudes and transients vary with the imposed parameters. Units are Hz, rad, and poise.

S11 Fig. Out-of-phase schooling angle across kinematic and viscosity conditions. Time histories of stagger angle *α* for Φ = *π*, corresponding to the conditions in Figure S10. The curves illustrate fluctuations about the predominantly side-by-side schooling geometry. Units are Hz, rad, and poise.

S12 Fig. In-phase schooling distance across kinematic and viscosity conditions. Time histories of relative distance *r* for Φ = 0 under the indicated pitching frequencies *f* , amplitudes Θ, and viscosities *µ*. Transients and oscillations vary across the conditions. Units are Hz, rad, and poise.

S13 Fig. In-phase schooling angle across kinematic and viscosity conditions. Time histories of stagger angle *α* for Φ = 0, corresponding to the conditions in Figure S12. The panels show parameter-dependent oscillations in the staggered schooling geometry. Units are Hz, rad, and poise.

S14 Fig. Foil trajectories during embodied schooling. (A) Out-of-phase (Φ = *π*) and (B) in-phase (Φ = 0) conditions in streamwise and lateral coordinates. The out-of-phase pair primarily advances streamwise, whereas the in-phase pair exhibits more pronounced lateral translation.

S15 Fig. Relationship between swimming number and Reynolds number in CFD and flow-tank experiments. Circles and squares denote in-phase and out-of-phase CFD cases; the dashed line shows the relationship plotted for the numerical results. Experimental cases (triangles) span Reynolds numbers near 10,800 and 15,000 and swimming numbers ^39^ of 12,000-43,200, placing the flow-tank conditions in context with the simulations.

S16 Fig. Flow-tank apparatus for the two-foil experiments. The setup comprises a recirculating flow tank, two tethered foils produced using a Bambu X1 3D printer, DYNAMIXEL XL430 pitching motors, and ME K6D40 six-axis sensors measuring streamwise and lateral forces.

S17 Fig. Force-balanced schooling candidates for out-of-phase foils (Φ = *π*). The maps show measured cycle-averaged force-difference vectors as functions of relative distance and stagger angle under the conditions specified in each panel. Arrows indicate force-difference directions, and colors indicate magnitudes. Marked candidate locations satisfy the local force-balance and directional criteria of Algorithm S2; they do not independently establish dynamic stability.

S18 Fig. Force-balanced schooling candidates for in-phase foils (Φ = 0). Measured cycle-averaged force-difference maps are shown against relative distance and stagger angle for the indicated conditions. Marked candidate staggered configurations are identified using Algorithm S2. These are constrained-foil force measurements, not free-swimming trajectories.

S19 Fig. Wake observations and idealization used in the Vortex Balance Model. (A,B) CFD wake structures for out-of-phase and in-phase foils. (C,D) Corresponding planar PIV wake visualizations. (E,F) Idealized vortex arrangements and induced-velocity components for the two phase conditions. The VBM represents selected wake interactions rather than the full three-dimensional flow.

S20 Fig. Definition of the effective tailbeat amplitude. (A,B) Transverse tailbeat trajectories in one-degree-of-freedom and two-degree-of-freedom motion, respectively. Translation of the foil modifies the actual amplitude relative to that inferred from prescribed pitching alone.

S21 Fig. Comparison between the present VBM predictions and previously reported schooling configurations. Symbols represent data reproduced from previous studies, including Newbolt et al.^34^, Ormonde et al.^35^, Peng et al. ^61^, Heydari et al.^62^, Lin et al.^63^ and Wang et al.^64^. The inset provides a separate scatter plot for the cases with *α* = *π/*2. Compared with previous studies of one-degree-of-freedom (1-DOF) systems, in which multiple schooling patterns emerged depending largely on the initial separation between swimmers, our 2-DOF simulations converged toward a characteristic time-averaged schooling configuration for each phase condition, across the tested initial separations (Figures S8–S13). Larger initial separations increased the time required for schooling to emerge (Figure S9). This difference suggests that allowing swimmers to move laterally, rather than restricting them to forward motion, can influence how schooling patterns emerge through hydrodynamic interactions. Thus, the freedom of movement available to individuals may be important in determining their collective organization.

S22 Fig. Randomized fish-pair control under combined visual and lateral-line impairment (V-LL-). (A) Relative configurations obtained by pairing individual trajectories from separate recordings while preserving their kinematics. (B) Corresponding VBM predictions. The comparison illustrates the reduced correspondence between randomly paired trajectories and the model.

S23 Fig. Repeatability of flow-tank force measurements. Representative (A) streamwise and (B) crossstreamwise force histories from three repetitions under identical conditions. The periodic signals show comparable amplitudes and phases across trials.

S24 Fig. Experimental test of lateral boundary sensitivity. Time histories of (A,B) streamwise and (C,D) crossstreamwise forces on foils #1 and #2 at a reference position and after shifting the pair toward the side wall. Similar signals indicate limited sensitivity to the tested lateral displacement, without independently quantifying free-surface effects.

S25 Fig. CFD assessment of domain-size effects on hydrodynamic forces. (A,B) Domains matching the experimental tank dimensions (8.75*BL ×* 2.5*BL*) and a larger domain (43.75*BL ×* 12.5*BL*), respectively. (C,D) Viscous force coefficients on foils #1 and #2; (E,F) corresponding pressure force coefficients. Reported relative differences are 3.34% and 3.41% for viscous contributions and 2.49% and 2.05% for pressure contributions. The comparison indicates limited domain-size sensitivity under the tested conditions; the calculation basis for these percentages should be specified.

S26 Fig. Experimental arrangement for tracking fish pairs. The diagram indicates the flow direction, coordinate axes, and synchronized side and reflected bottom views. A mirror at 45^°^ provides the bottom view for reconstructing three-dimensional relative positions.

S27 Fig. Tailbeat-frequency distributions under the four sensory conditions. Frequency distributions for the two fish in each pair are shown for intact vision and lateral line (V+LL+), visual deprivation (V-LL+), lateral-line impairment (V+LL-), and combined impairment (V-LL-). Tailbeat frequencies were obtained from tracked body undulations using wavelet analysis.

S28 Fig. Tailbeat-amplitude distributions under the four sensory conditions. Amplitudes for the two fish are shown under V+LL+, V-LL+, V+LL-, and V-LL-conditions. Amplitudes were extracted from reconstructed body motion; these kinematic measurements were used as inputs to the VBM.

S29 Fig. Fish-tracking workflow for extracting swimming kinematics. Mask R-CNN segments each fish silhouette, and DeepPoseKit identifies ordered landmarks to reconstruct the body centerline. The resulting motion records provide body undulation, tailbeat amplitude, phase, and frequency for the biological comparison with the VBM.

S1 Text Numerical methodology

S2 Text Flow-tank experiments and boundary effects S3 Text Vortex Balance Model

S4 Text Analysis of CFD trajectories

S5 Text Particle image velocimetry measurements S6 Text Fish tracking and statistical analysis

S1 Video. CFD trajectories illustrating the emergence of embodied schooling under prescribed foil pitching.

S2 Video. Examples of randomized CFD trajectory pairs used to assess coincidental schooling-like motion.

S3 Video. Flow-tank measurements of the hydrodynamic forces acting on two pitching foils at prescribed relative positions.

S4 Video. Comparison between fish movement and VBM predictions.

S5 Video. CFD flow-field visualizations showing the vortex structures associated with the two phase conditions.

S1 Algorithm. Detecting embodied schooling and determining the final relative configuration of two numerical foils

S2 Algorithm. Detecting a convergent zero-force point from the measured force-vector field

## Acknowledgments

We would like to express our gratitude to Dr Z.S., Y.Z., and Mr P.Y., R.W. at Max Planck Institute of Animal Behaviour for sharing their computational resources. We acknowledge funding support from the Max Planck Society, the National Natural Science Foundation of China 12502279 (L.-M.C.), the Office of Naval Research N00014-64019-1-2556 (I.D.C.), the European Union’s Horizon 2020 research and innovation programme under the Marie Sklodowska-Curie grant agreement 860949 (I.D.C.), the PathFinder European Innovation Council Work Programme 101098722 (I.D.C.), the Struktur-und Innovationsfonds für die Forschung (SI-BW) of the State of Baden-Württemberg, the Deutsche Forschungsgemeinschaft (DFG, German Research Foundation) under Germany’s Excellence Strategy-EXC 2117–422037984, DFG project 462886202 (I.D.C.), the Deutsche Forschungsgemeinschaft Gottfried Wilhelm Leibniz Prize 2022 584/22 (I.D.C.), the Sino-German Centre in Beijing for generous funding of the Sino-German mobility grant M-0541 (L.L.), Messmer Foundation Research Award (L.L.), National Science Foundation Awards OAC 1931516 (B.E.G.), National Science Foundation Awards OAC 1931368 and CBET 2234387 (A.P.S.B.).

## Author contributions

L.-M.C. and L.L. conceived the idea and designed the project. L.-M.C. carried out the CFD simulations. L.-M.C., H.J. and L.L. performed the flow tank experiments. L.L. conducted the biological experiments. L.-M.C. and L.J. formulated the Vortex Balance Model (VBM). L.-M.C., L.L., and M.N. conducted data analysis. A.P.S.B. and B.E.G. provided the numerical simulation platform and technical support. I.D.C. provided resources, feedback on experimental design, and guidance throughout the project. L.-M.C. and L.L. wrote the initial draft with input from I.D.C., and all authors contributed to manuscript revision and interpretation of results.

